# Chimeric Antigen Receptor Macrophages Target and Resorb Amyloid Plaques in a Mouse Model of Alzheimer’s Disease

**DOI:** 10.1101/2023.04.28.538637

**Authors:** Qiuyun Pan, Ping Yan, Alexander B. Kim, Qingli Xiao, Gaurav Pandey, Hans Haecker, Slava Epelman, Abhinav Diwan, Jin-Moo Lee, Carl J. DeSelm

## Abstract

Substantial evidence suggests a role for immunotherapy in treating Alzheimer’s disease (AD). Several monoclonal antibodies targeting aggregated forms of beta amyloid (Aβ), have been shown to reduce amyloid plaques and in some cases, mitigate cognitive decline in early-stage AD patients. We sought to determine if genetically engineered macrophages could improve the targeting and degradation of amyloid plaques. Chimeric antigen receptor macrophages (CAR-Ms), which show promise as a cancer treatment, are an appealing strategy to enhance target recognition and phagocytosis of amyloid plaques in AD. We genetically engineered macrophages to express a CAR containing the anti-amyloid antibody aducanumab as the external domain and the Fc receptor signaling domain internally. CAR-Ms recognize and degrade Aβ *in vitro* and on APP/PS1 brain slices *ex vivo;* however, when injected intrahippocampally, these first-generation CAR-Ms have limited persistence and fail to reduce plaque load. We overcame this limitation by creating CAR-Ms that secrete M-CSF and self-maintain without exogenous cytokines. These CAR-Ms have greater survival in the brain niche, and significantly reduce plaque load locally *in vivo*. These proof-of-principle studies demonstrate that CAR-Ms, previously only applied to cancer, may be utilized to target and degrade unwanted materials, such as amyloid plaques in the brains of AD mice.

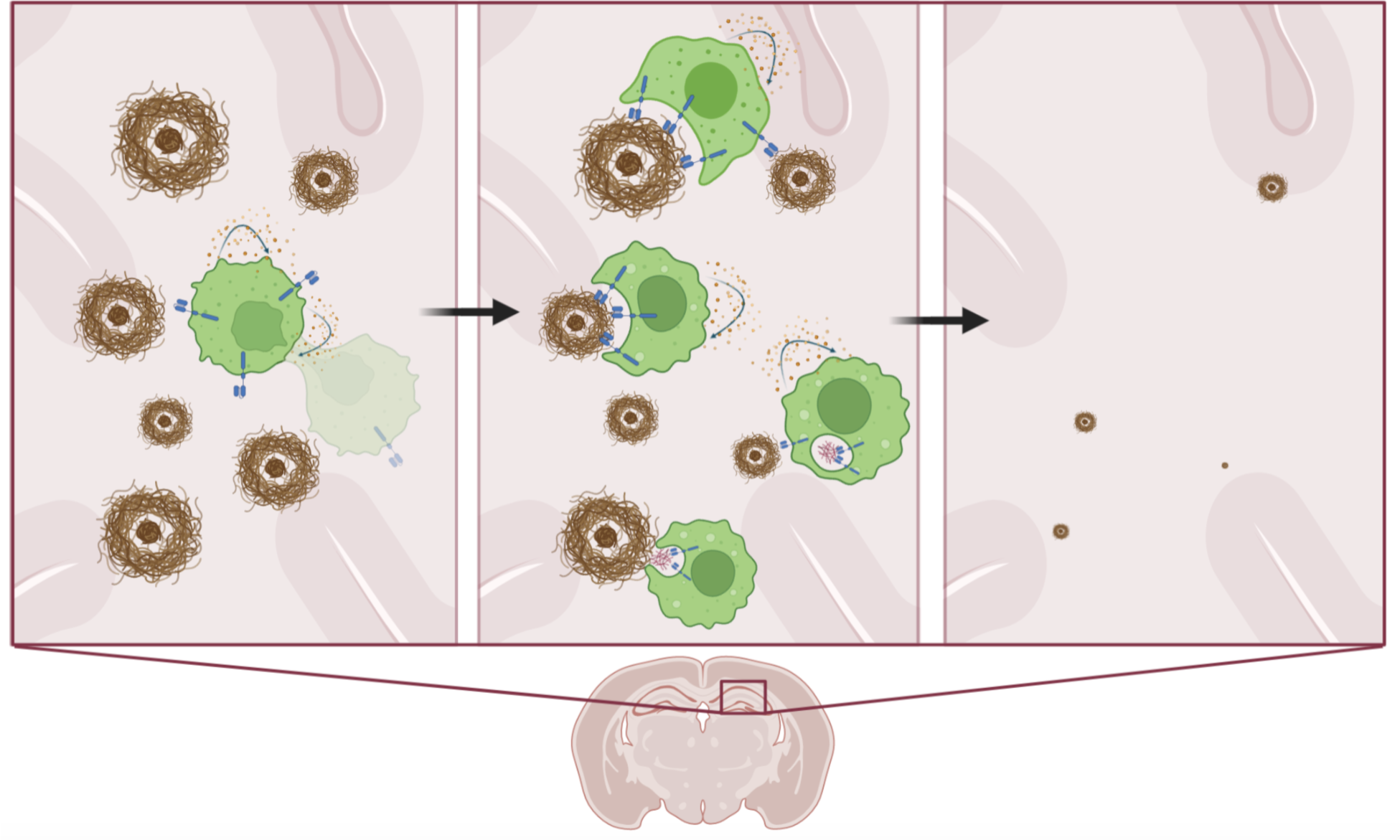
Amyloid targeting CAR Macrophages engineered to secrete M-CSF promote their own local survival and expansion while resorbing amyloid plaques in the brains of Alzheimer’s disease APP/PS1 mice, resulting in significant local clearance of amyloid plaques of all sizes.

## Introduction

Recent breakthroughs in liquid cancer treatment brought on by the advent of CAR T-cells (Mailankody et al., 2022; Neelapu et al., 2017; Park et al., 2018; Raje et al., 2019; Wang et al., 2020) inspire considerations that CAR cellular therapy, in which a specialized immune cell is modified with a chimeric antigen receptor (CAR) to both target a specific antigen and induce a desired cellular function in response to that antigen, may have broader roles in medicine beyond cancer. Indeed, the idea of a living drug that can respond in more sophisticated ways than a single small molecule is appealing for diseases that have thus far evaded conventional approaches. Beyond cancer, Alzheimer’s disease (AD) is one such disease that is both growing in prevalence and partially treatable but not curable, despite a large number of mouse and human trials with small molecules, antibodies, and targeted therapies (Karran and De Strooper, 2022; van Dyck et al., 2023). The pathophysiology of AD is still incompletely understood; however, beta amyloid (Aβ) plaque deposition is thought to be a key initial trigger for neurodegeneration (Scheltens et al., 2021). Antibodies to Aβ have recently been shown to reduce amyloid plaque load and mitigate cognitive decline in AD patients (Budd Haeberlein et al., 2022; van Dyck et al., 2023), but also have dose-limiting side effects such as amyloid-related imaging abnormalities (ARIA) (Salloway et al., 2022), including hemorrhage and edema ranging from non-symptomatic to lethal (Couzin-Frankel and Piller, 2022; Reish et al., 2023), without fully stopping its cognitive decline. New classes of therapies in this area may herald new benefits.

Previously, CAR-Macrophages (CAR-Ms) have been reported to phagocytose tumor cells (Klichinsky et al., 2020), and are currently in a cancer clinical trial (NCT04660929). CAR-M technology has thus far not been extended to non-cancerous conditions. Microglia, the resident immune cells in the brain, phagocytose and degrade Aβ and amyloid plaques in AD, but are noted to be dysfunctional with amyloid engorgement and dispensable for progression of plaque pathology. Ontogenically distinct circulating peripheral monocytes of adult or definitive hematopoietic origin are recruited to amyloid plaques in the APP/PS1 transgenic mouse model of AD, where they constitute 6% of all plaque-associated macrophages and function in attenuating amyloid plaque load (Yan et al., 2022). Enhancing their capacity to resorb plaque may yield further benefits.

We wondered if introducing a CAR into macrophages that targets Aβ on the extracellular portion (using the aducanumab single-chain variable fragment, scFv), with the phagocytic common gamma chain of the Fc receptor on the intracellular portion, would yield effective Aβ phagocytosis. The ultimate goal is to create a conceptually new form of therapy for AD with potentially favorable risks and benefits. We find that this first-generation CAR-M resorbs plaque *in vitro* and *ex vivo*, but has limited *in vivo* survival and fails to reduce local plaque load *in vivo*. However, a next-generation, self-sustaining CAR-M that secretes M-CSF was shown to expand in the brain microenvironment after four days of microglia depletion with the CSF-1 inhibitor PLX5622. Furthermore, these cells significantly resorbed plaques locally in the hippocampi of aged APP/PS1 mice compared to control M-CSF secreting non-targeting CAR-Ms. These studies demonstrate that targeted CAR-Ms may be utilized to resorb Aβ *in vivo*, and suggest living drugs may be considered in potential future treatment approaches for AD.

## Results

### Aβ CAR-Ms significantly bind and resorb beta-amyloid in culture and on brain slices *ex vivo*

Since this is a novel CAR construct (Fig. 1a), we first tested whether it is successfully expressed on the surface of macrophages after transduction and differentiation. Using an antibody that directly recognizes the CAR, we find high surface expression can be obtained on transduced HoxB8 cells (Fig. S1a), which were generated by transducing bone marrow cells with the previously described estrogen-responsive HoxB8 construct. This construct sustains the cells in an undifferentiated state in the presence of estrogen while maintaining the ability to normally differentiate into macrophages in the presence of M-CSF without estrogen (Redecke et al., 2013). CAR-expressing HoxB8 cells were sorted, and the expression of CAR was confirmed on HoxB8 cells differentiated into mature macrophages (defined by F4/80 and CD64) with M-CSF for 6 days (Fig. 1b). To test whether the scFv expressed on the surface of M-CSF differentiated CAR-Ms maintains the ability to bind Aβ, Aβ CAR-Ms or control CAR-Ms, which target an irrelevant antigen EphA2 and lack an intracellular domain (Fig.1a), were co-cultured with HiLyte™ Fluor 488-labeled Aβ(1-42) for different time intervals. Cells were washed and analyzed by flow cytometry, which demonstrated a CAR-dependent two-fold increase in both the percentage of cells that had taken up Aβ as well a significant increase in the amount of Aβ taken up by each cell, both increasing in a time-dependent fashion (Fig. 1c). Increased Aβ uptake by Aβ CAR-Ms over control CAR-Ms could also be visualized by microscopy (Fig. S1b), which confirmed cellular binding of Aβ(1-42) (Fig. 1d).

**Figure 1:**
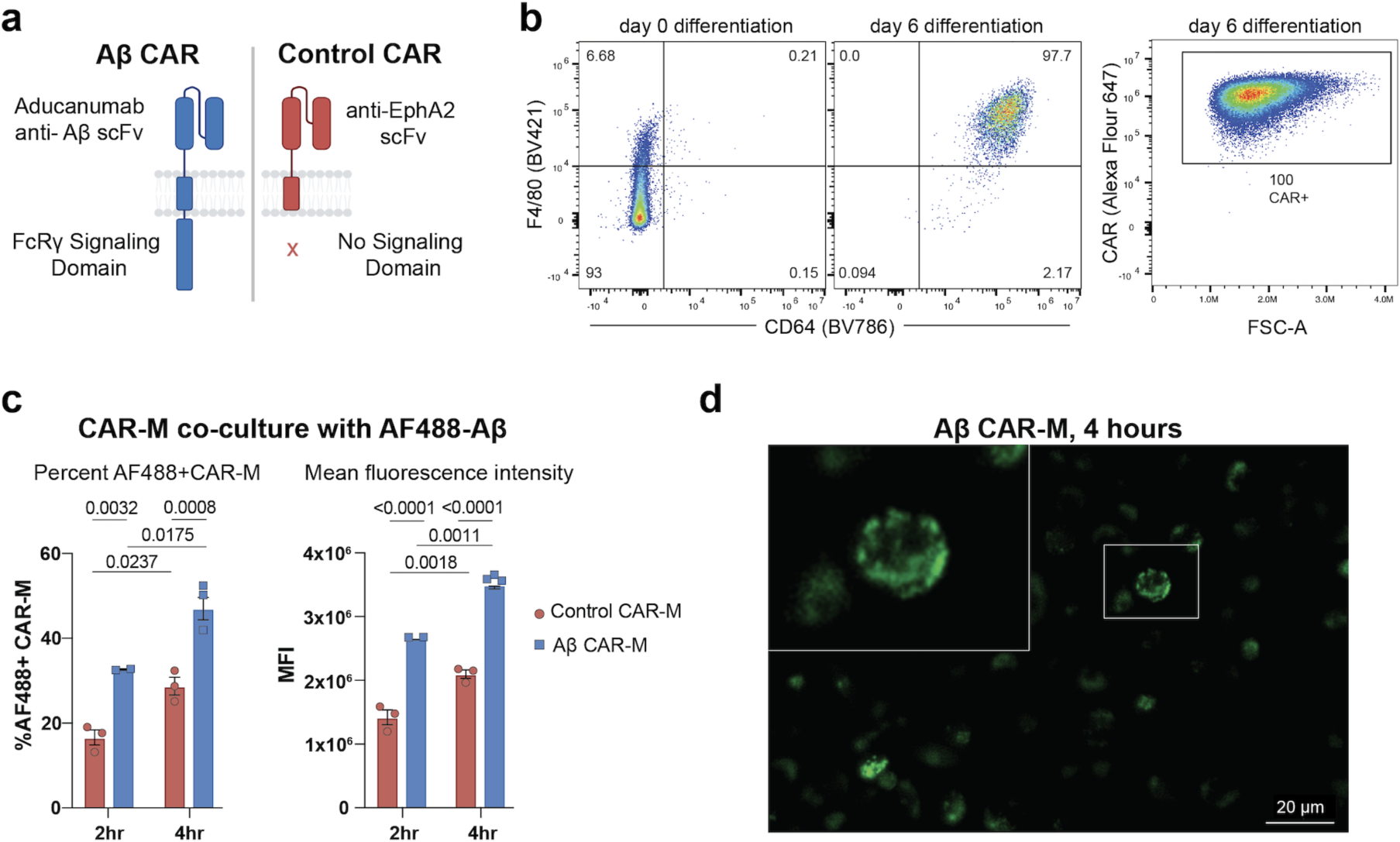
Generation of an aducanumab-based Aβ CAR-M and validation of CAR-mediated uptake of soluble Aβ. **a)** Schematic diagram of Aβ CAR and Control CAR constructs. **b)** Representative FACS plots of (left) F4/80 and CD64 surface expression on HoxB8 cells at day 0 and day 6 of differentiation with M-CSF; (right) surface CAR expression on cells at day 6 of differentiation. FSC-A, forward scatter area. **c)** *In vitro* phagocytosis of AlexaFlour488 fluorescent tagged Aβ(1-42) by Aβ CAR Ms or control CAR-Ms after 2 or 4 hours of co-incubation, depicted as % uptake (left) or mean fluorescence intensity (MFI, right). Data is represented as mean ± s.e.m. from n=2-3 independent experiments with 2-3 technical replicates for each condition. Statistical significance was calculated with 2-way ANOVA with *Šidàk*’s multiple comparisons test **d)** Representative image of Aβ CAR Ms co-cultured with AF488 tagged Aβ(1-42) for 4 hours.

To assess a more physiologically relevant target, we cultured Aβ-targeted or control CAR-Ms on *ex vivo* brain slices from aged APP/PS1 mice, to assess their ability to resorb plaques that were deposited *in vivo* (Fig. 2a). Control and Aβ CAR-Ms were cultured on adjacent brain slices to approximately match plaque load and distribution. Quantification of HJ3.4-stained plaques, which recognizes all forms of Aβ (Funk et al., 2015), revealed a significant reduction in plaque burden in Aβ CAR-M treated slices over control CAR-M slices, which on their own exhibit some resorptive activity, *ex vivo* (Fig. 2b). In preparation for *in vivo* studies, we desired the ability to non-invasively image the CAR-Ms over time, so we retrovirally introduced GFP-Luciferase into Aβ and control CAR-Ms. To assess whether expressing GFP-Luciferase (GFP-Luc) affects CAR-M function, we repeated *ex vivo* plaque resorption assays comparing these cells. To further differentiate resorption of all Aβ species from fibrillar compact plaques, we stained slices with HJ3.4 antibody as well as X-34, respectively (Styren et al., 2000). We found that Aβ CAR-Ms could effectively reduce both compact plaques (X-34-stained) as well as diffuse plaques of all Aβ species (HJ3.4-stained). This activity was observed throughout the whole brain slice, regardless of GFP-Luc expression (Fig. 2c, 2f). Thus, subsequent studies used GFP-Luc expressing control and Aβ CAR-Ms.

**Figure 2:**
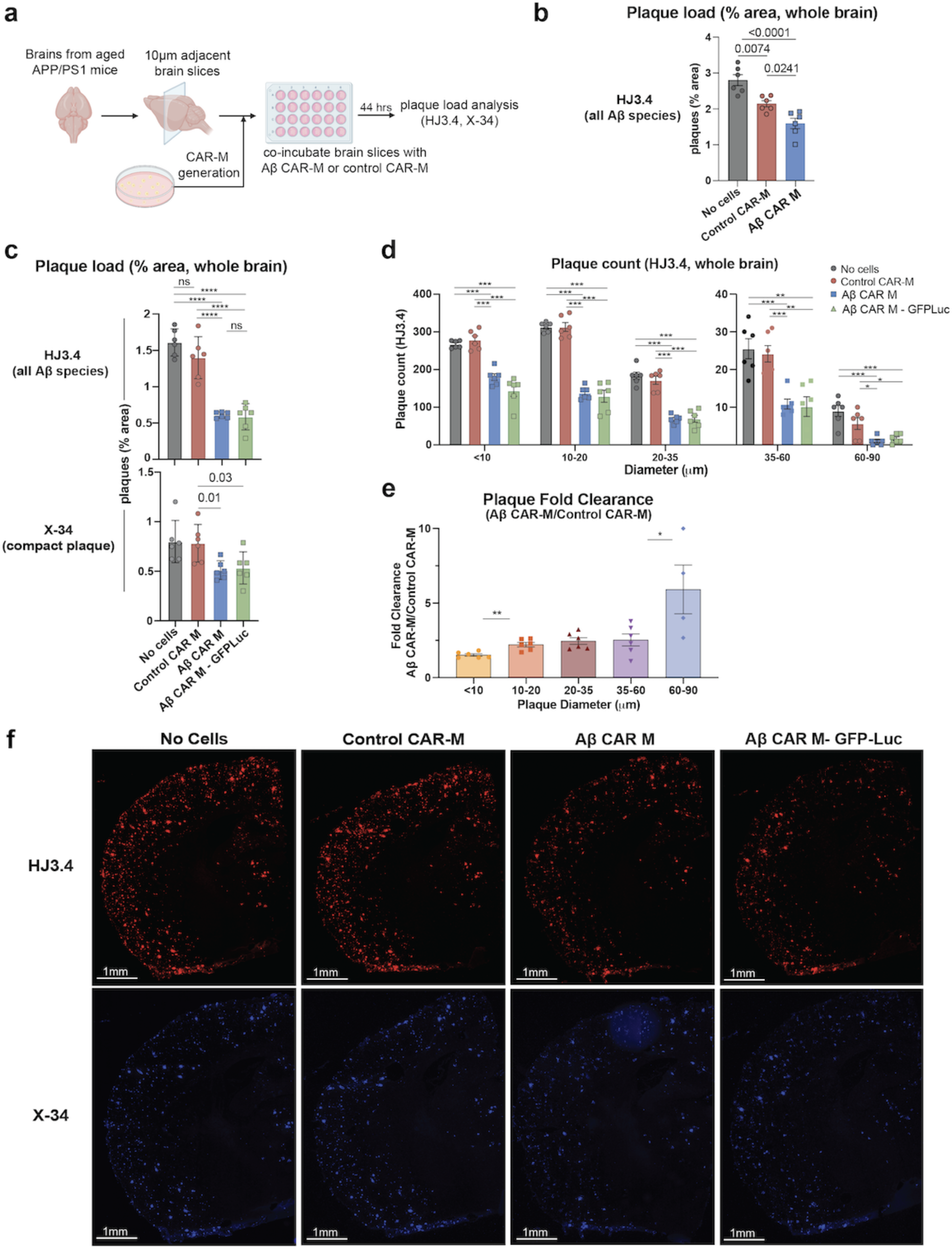
Aβ CAR-M resorb amyloid plaques of various sizes on brain slices from aged APP/PS1 mice *ex vivo*. **a)** Schematic of *ex vivo* assessment of amyloid plaque phagocytosis. Adjacent brain slices from aged APP/PS1 mice were co-incubated with Aβ CAR-M or Control CAR-Ms for 44 hours and plaque load was assessed with HJ3.4 or X-34 immunostaining. **b-e)** Assessment of plaque load **(b, c)**, plaque count **(d)**, and plaque fold clearance of Aβ CAR-M over Control CAR-Ms **(e)** on APP/PS1 brain slices after co-incubation with no cells, control CAR-M, or Aβ CAR-M with or without GFP-Luc. Data shown as mean ± s.e.m from n=5-6 independent experiments with 5-6 technical replicates each. Statistical significance was calculated with one-way ANOVA with Tukey’s multiple comparisons test (**c (**HJ3.4**), d**) or unpaired t-tests (**c** (X-34), **e**). For **c-e**, *P < 0.05, **P < 0.01, ***P < 0.001, ****P < 0.0001. ns, not significant. **f)** Representative images demonstrating adjacent brain sections stained with HJ3.4 or X-34 after co-incubation with no cells, control CAR-M, Aβ CAR-M, or Aβ CAR-M GFP-Luc cells.

### CAR-Ms resorb *ex vivo* amyloid plaques of all sizes

Macrophages are known to phagocytose particles most effectively in the 1-3 micron size range (Champion et al., 2008; Pratten and Lloyd, 1986; Tabata and Ikada, 1988), with a rapid fall-off in phagocytic capacity of particles of >3 microns in diameter (Pacheco et al., 2013). Although the average plaque size is much larger than this – most commonly 10-20 microns – Aβ CAR-Ms effectively resorb plaque of all sizes (Fig. 2d). In fact, when we assessed fold change in plaque clearance of Aβ-targeted versus control CAR-Ms, targeted CAR-Ms resorbed larger plaques of >10 microns relatively more effectively than smaller plaques (Fig. 2e). This, as well as direct immunohistological visualization of plaque-CAR-M interactions (Fig. 3f, S2e), suggests that plaque is likely resorbed in pieces rather than being taken up by whole plaque phagocytosis.

**Figure 3:**
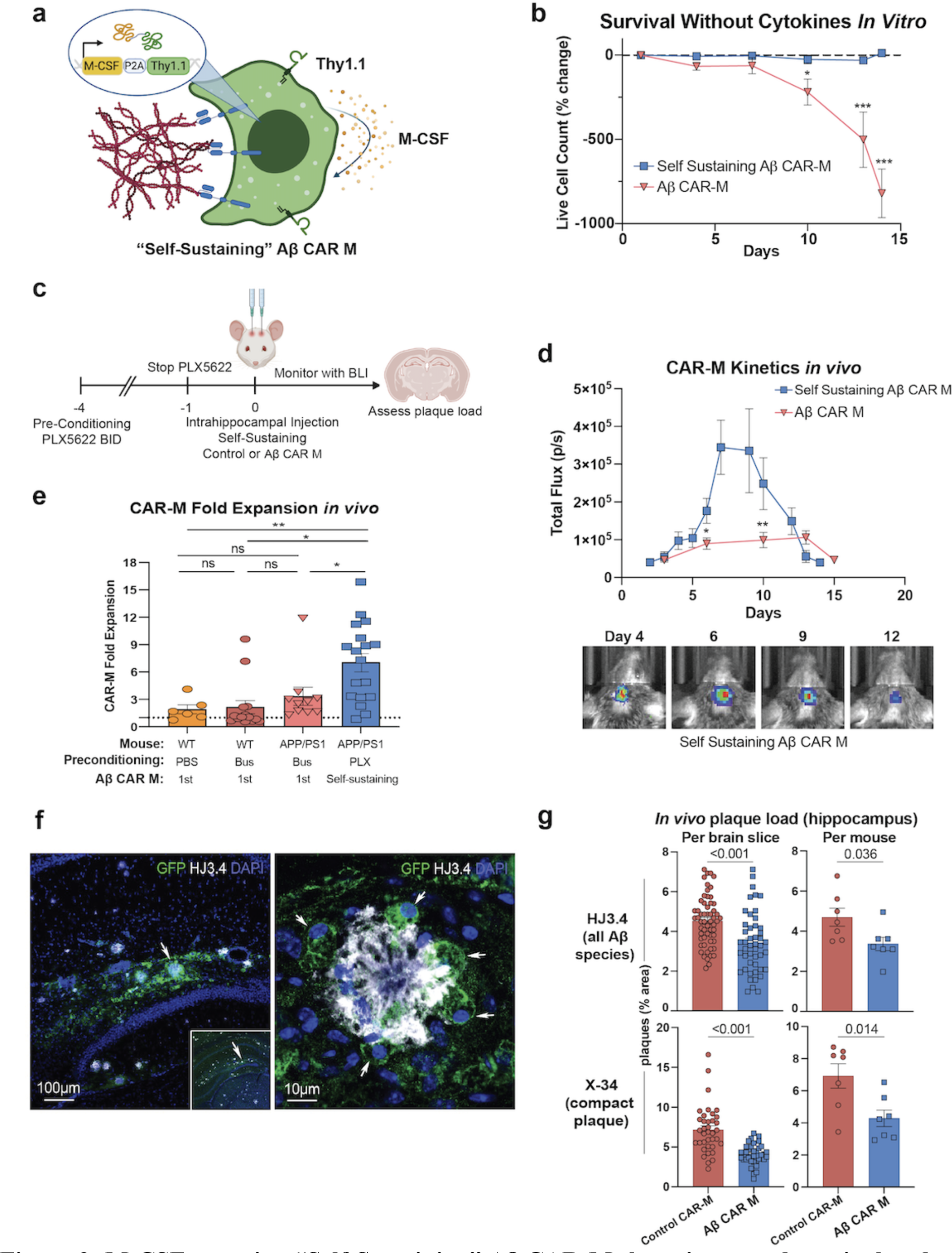
M-CSF secreting “Self-Sustaining” Aβ CAR-Ms have improved survival and reduce plaque load *in vivo*. **a)** Schematic of M-CSF secreting “self-sustaining” construct retrovirally introduced into control and Aβ CAR-Ms that contains the M-CSF gene followed by a P2A cleavage sequence and Thy1.1. **b)** % change in live cell count of first-generation Aβ CAR-Ms and self-sustaining Aβ CAR-Ms upon removal of M-CSF from the culture medium *in vitro*, determined by flow cytometry staining with ZombieNIR live/dead staining. Cells were differentiated for 6 days in M-CSF to become mature macrophages, prior to M-CSF removal. Statistical significance was calculated with an unpaired t-test. **c)** Schematic of PLX5622 pre-conditioning and intrahippocampal injection of self-sustaining Aβ CAR-Ms. **d)** Total flux determined by non-invasive bioluminescence imaging (BLI) tracking first-generation Aβ CAR-M kinetics *in vivo* compared to self-sustaining Aβ CAR-M kinetics after intrahippocampal injection (upper). Representative BLI images from self-sustaining Aβ CAR-M treated mice (lower). Days indicates days post-intrahippocampal injection. n=10-18 mice per group. Statistical significance was calculated with unpaired t-tests. **e)** Fold-expansion of CAR-Ms from the first day of BLI after intrahippocampal injection of cells to the day of maximum total flux measured by BLI. n=6-18 mice per group. Statistical significance was calculated with one-way ANOVA with Tukey’s multiple comparisons test. **f)** Representative immunofluorescence microscopy images of self-sustaining Aβ CAR-Ms binding to amyloid plaque *in vivo*. **g)** Assessment of plaque load after intrahippocampal injection of self-sustaining control CAR-M or self-sustaining Aβ CAR-M in n=7 aged APP/PS1 mice. Mice were sacrificed on day 12 or 13 post intrahippocampal injection and brain tissue was sectioned and stained with HJ3.4 and X-34 to assess plaque load. Data shown as mean ± s.e.m. Statistical significance was calculated with unpaired t-tests. For **b-d**, *P < 0.05, **P < 0.01, ***P < 0.001, ****P < 0.0001, ns, not significant.

### First-generation Aβ CAR-Ms find plaque *in vivo* but have limited expansion and survival

While *ex vivo* assays demonstrated the proof of concept that Aβ CAR-Ms can specifically bind and resorb plaque on two-dimensional brain slices, *in vivo* function relies on the ability of these peripherally-derived cells to adapt to the brain niche, compete with local resident cell types, and find and resorb plaques in three-dimensional space. Given the importance of preconditioning in CAR T-cell therapy (Gattinoni et al., 2005; Giuffrida et al., 2020; Murad et al., 2021) as well as in prior microglia engraftment studies (Xu et al., 2020), we preconditioned mice with low dose (non-ablative) Busulfan (Wilkinson et al., 2013; Youshani et al., 2019) for two days to deplete endogenous microglia prior to intrahippocampal injection of PBS or Aβ CAR-Ms in PBS with M-CSF (Fig. S2a). Because it was uncertain if the CAR-Ms would spread within the brain after intrahippocampal injection, we chose to inject one hippocampus with cells and the contralateral hippocampus with PBS. We assessed the overall engraftment of CAR-Ms using bioluminescence imaging in both WT and APP/PS1 mice, and found that in APP/PS1 mice, CAR-Ms survived and modestly expanded for the first 10 to 12 days, but subsequently rapidly diminished to baseline by day 14 (Fig. S2b, S2c). Mice were sacrificed on day 14 and plaque load was quantified in a circular region of interest (ROI) surrounding the PBS or Aβ CAR-M injection site. This circular ROI was selected based on the cross-sectional area of the injection site and was centered around residual GFP signal that was visible in the hippocampal region (Fig. S2d).

Uniform circle sizes were used across all slices, and as a control, random areas from remote sites of the cortex were analyzed to ensure the density of plaques was relatively uniform between slices/mice. A small number of Aβ CAR-Ms remained, and some were observed to be recognizing and binding to Aβ plaques (Fig. S2e), however no significant difference was observed in overall plaque load between PBS and Aβ CAR-M treated hemispheres, even surrounding the local site of injection (Fig. S2f). Because the injected CAR-Ms appeared to stay relatively localized to the injection site (Fig. S2d), for subsequent studies we decided to inject the left hippocampus of each animal with control CAR-Ms and the right hippocampus of each animal with Aβ CAR-Ms. This bilateral injection paradigm was designed to control for inter-mouse heterogeneity in hippocampal plaque load.

### M-CSF expressing CAR-Ms have enhanced survival and locally reduce hippocampal plaque load *in vivo*

Due to the lack of survival, persistence, and efficacy of first-generation CAR-Ms *in vivo*, we developed M-CSF-expressing CAR-Ms to enhance cell-autonomous differentiation and survival and improve persistence and efficacy *in vivo*. We co-infected control or Aβ CAR-Ms with a retroviral construct containing M-CSF followed by a P2A cleavage sequence and Thy1.1 for sorting (Fig. 3a). Since Thy1.1 is expressed at a 1:1 ratio with M-CSF, we sorted cells of the same Thy1.1 mean fluorescence intensity (MFI) to obtain consistently and uniformly expressed M-CSF by both control and Aβ CAR-Ms (Fig. S3a). While first-generation CAR-Ms steadily and nearly completely die within a two-week period without exogenous cytokine, we found that M-CSF-expressing CAR-Ms were able to persist at a stable level *in vitro* without significant death or expansion in the absence of any exogenous cytokine (Fig. 3b). Since these M-CSF-secreting CAR-Ms maintain completely independently of cytokine support, we refer to them as self-sustaining CAR-Ms.

To assess *in vivo* survival and expansion of self-sustaining CAR-Ms in the brain niche, we injected self-sustaining control and Aβ CAR-Ms in opposite hippocampi of aged APP/PS1 mice. To deplete endogenous microglia while sparing non-myeloid cells with a potentially more clinically translatable drug (Basilico et al., 2022; Spangenberg et al., 2019), we subjected mice to four days of preconditioning with the selective CSF-1 inhibitor PLX5622 (Fig. 3c). Bioluminescence imaging showed self-sustaining CAR-Ms rapidly expanded in the first week within the brain, then plateaued and rapidly contracted (Fig. 3d, S3b). Although we saw only modest microglial depletion with this preconditioning regimen compared with PBS control or with low-dose Busulfan preconditioning (Fig. S3c), self-sustaining CAR-Ms reached significantly greater numbers than first-generation CAR-Ms, as quantified by non-invasive bioluminescence imaging over time (Fig. 3d). To assess whether CAR-Ms were merely persisting or were actually expanding after *in vivo* injection in our various models, we quantified fold change in bioluminescence total flux at peak CAR-M levels relative to baseline at injection. We found that, on average, little expansion occurred with first-generation CAR-Ms in any of our models, and in several mice, the maximum CAR-M load occurred at baseline after injection, indicating little to no proliferation *in vivo* (Fig. 3e). However, self-sustaining CAR-Ms expanded in all but one of 18 injected mice, with an average 7-fold expansion (Fig. 3e).

Histologic analysis of brain slices from treated mice demonstrated that Aβ CAR-Ms again recognized plaque *in vivo* and demonstrated plaque phagocytosis (Fig. 3f). Furthermore, unlike prior experiments using first-generation CAR-Ms, quantification of plaque load in the same ROI as defined in prior studies around the hippocampal injection site (Fig. S3d) revealed significant plaque reduction after self-sustaining Aβ CAR-M treatment compared to self-sustaining control CAR-M treatment in the contralateral hippocampus (Fig. 3g). The reduction in Aβ plaque load was evident with self-sustaining Aβ CAR-Ms regardless of whether individual brain slices were analyzed, or average plaque load per mouse was analyzed (Fig. 3g). Additionally, compact plaques, stained with X-34, were also significantly reduced by self-sustaining Aβ CAR-Ms (Fig. 3g). Very little migration of CAR-Ms was observed from the site of injection (Fig. S2d), and therefore, plaque reduction was only observed within the vicinity of CAR-M injection. Indeed, analysis of random remote regions of interest of identical volume in mirror regions of the cortex did not reveal any changes in plaque load (Fig. S3e), demonstrating relative uniformity of plaque load across hemispheres. While these results demonstrate proof of principle that CAR-Ms can focally reduce amyloid plaque load, further work is needed to generate CAR-Ms capable of broad migration to reduce plaque load throughout the brain.

## Discussion

In this study, we tested the ability of macrophages expressing a novel Aβ-targeting chimeric antigen receptor (CAR-Ms) to bind and phagocytose Aβ and amyloid plaques. We found that Aβ-targeting CAR-Ms uptake Aβ peptide *in vitro* in a CAR-dependent manner, and significantly reduce both Aβ plaque load and compact plaque load when cultured on brain slices from aged APP/PS1 mice *ex vivo*. We further show that Aβ-targeting CAR-Ms engineered to self-sustain by producing M-CSF significantly reduce hippocampal Aβ plaque load and compact plaque load *in vivo* when injected into the hippocampus of aged APP/PS1 mice after preconditioning with the CSF-1 inhibitor PLX5622 to deplete endogenous microglia. Together, our findings support the further development of CAR-Ms as a potential anti-Aβ therapeutic approach for treating Alzheimer’s disease (AD).

These results demonstrate that while macrophages expressing a CAR targeting Aβ can significantly resorb Aβ and amyloid plaques *in vitro* and *ex vivo*, achieving significant *in vivo* plaque resorption required engineering the cells to self-sustain by producing M-CSF for autocrine and/or paracrine stimulation. While M-CSF secretion alone may have achieved some degree of plaque clearance through either microglia stimulation (Boissonneault et al., 2009) or improving CSF flow dynamics (Drieu et al., 2022), we measured a significant independent CAR target-mediated effect, since our control hemispheres were also treated with M-CSF secreting CAR-Ms but not targeting Aβ.

In several neurological diseases, including AD, the microglial niche is known to undergo phenotypic changes that can be both beneficial and detrimental to disease progression (Salter and Stevens, 2017). Because of these insights, there have been several efforts to replace dysfunctional microglia with healthy microglia or with peripheral monocytes via microglial or bone marrow transplant after preconditioning with myeloablative chemotherapy or radiation to remove endogenous microglia (Xu et al., 2020) (Shibuya et al., 2022). These studies also showed that while monocytes do not infiltrate the brain parenchyma under homeostatic conditions, monocytes can infiltrate the brain parenchyma in the setting of preconditioning chemotherapy or radiation, which in addition to depleting endogenous microglia to make room for new cells to engraft, presumably promotes peripheral monocyte infiltration into the brain due to disruption of entry sites into the central nervous system, including the blood-brain-barrier, and increased secretion of cytokines and chemokines that recruit peripheral immune cells (Ajami et al., 2007) (Marchetti and Engelhardt, 2020). In line with these studies, we also observed that preconditioning with agents that reduce endogenous microglia, in our case non-myeloablative (low dose) Busulfan (Yu et al., 2019), improves engraftment of peripheral myeloid cells in both WT and APP/PS1 mice compared to PBS preconditioning. Better microglial removal has been achieved with myeloablative doses of Busulfan followed by bone marrow transplant (Sailor et al., 2022); however, we intentionally did not pursue fully ablative regimens that have little potential for clinical translation. PLX5622 is reported to robustly deplete microglia as well as all other myeloid cells in other models if given ad libitum in the chow for an extended period (Henry et al., 2020; Spangenberg et al., 2019). In our studies, this more specific microglia-depleting preconditioning regimen was associated with significantly reduced plaque load in the hippocampus of Aβ CAR-M treated APP/PS1 mice. Since a similar PLX CSF1R inhibitor is currently FDA approved for a rare myeloid-derived tumor and has been well tolerated (Lamb, 2019; Tap et al., 2019), this may be a more clinically translatable preconditioning regimen. We appreciated only modest microglia reduction after delivering the drug IP BID for four days, and although microglia are expected to quickly recover after discontinuation of PLX5622, we still observed substantial proliferation of self-sustaining CAR-Ms in that setting.

It is thought that anti-Aβ monoclonal antibodies clear amyloid plaques by both cell-mediated and non-cellular mechanisms. Non-cellular mechanisms of amyloid plaque removal include direct dissolution of plaques and clearance of amyloid fibrils and oligomers by perivascular clearance pathways (Frenkel et al., 1999; Klyubin et al., 2005; Solomon et al., 1997). Another non-cellular mechanism of amyloid plaque removal is based on the peripheral sink hypothesis, which posits that peripherally administered antibodies shift the balance between CNS and peripheral Aβ by reducing peripheral Aβ, which promotes Aβ efflux from the brain to re-balance this equilibrium (DeMattos et al., 2001). A downside to non-cellular clearance is that the direct disaggregation of plaques and subsequent clearance of Aβ may promote cerebral amyloid angiopathy (Weller et al., 2009). Cell-mediated clearance of amyloid plaques by monoclonal antibodies is thought to occur in part by increasing the uptake of antibody coated plaque material by endogenous microglia through Fc receptor mediated phagocytosis (Bard et al., 2000; Schenk et al., 1999; Sevigny et al., 2016; Wilcock et al., 2004), though amyloid clearance has also been shown with F(ab’)2 antibody fragments lacking an Fc domain (Bacskai et al., 2001). These findings imply that both non-cellular and cell-mediated mechanisms play a role in the effectiveness of anti-Aβ monoclonal antibodies.

Accumulating evidence suggests that peripheral monocytes/macrophages phagocytose and degrade Aβ plaques more effectively than microglia. In one study, transgenic mice that block peripheral myeloid cell infiltration into the brain parenchyma have increased Aβ plaque load compared to control APP_Swe_/PS1 mice, suggesting that peripherally derived myeloid cells phagocytose and eliminate plaques better than brain-resident microglia (Simard et al., 2006). Conversely, the depletion of microglia in APP/PS1 mice led to no change in amyloid plaque count or size that developed over time, again implying that microglia are unable to effectively phagocytose and degrade amyloid plaques, though the ability of peripheral macrophages to reduce amyloid plaque load was not assessed in this study (Grathwohl et al., 2009). The inability of late stage disease microglia to control plaque load with AD progression may be attributed to the reduced expression of Aβ binding receptors and Aβ-degrading enzymes as plaque deposition increases with age, which was shown to lead to decreased phagocytosis and degradation of amyloid material in APP/PS1 mice (Hickman et al., 2008). Similarly, microglia isolated from human AD brains show reduced expression of molecules important for phagocytosis and the recycling of phagocytic receptors (Lucin et al., 2013). Recent reports suggest that microglia may even promote the spread and development of Aβ plaques (d’Errico et al., 2022; Huang et al., 2021). Peripheral macrophages, however, may also be defective at phagocytosing Aβ plaques in the setting of AD potentially due to chronic Aβ exposure, as it has been shown that monocytes/macrophages from AD patients were less effective at phagocytosing Aβ compared to cells from age-matched non-AD patients (Fiala et al., 2005). Allogenic CAR-M therapy may be a solution to overcome dysfunctional cell-mediated amyloid plaque clearance in AD and provides a theoretical advantage over therapeutic strategies that require functional endogenous cells to remove Aβ.

While we did not perform a direct comparison between Aβ CAR-Ms and anti-Aβ monoclonal antibodies, additional important theoretical advantages of using CAR-Ms to target amyloid plaques are that 1) they can be engineered to express additional helpful factors (such as enzymes that help degrade amyloid or other molecules, proteins that improve neuron or other functions, cytokines that may have independent benefits, etc.), 2) they may be polarized or engineered to acquire a desired phenotype, and 3) they constitutively phagocytose and degrade plaque material due to endogenous CAR expression without relying on antibody encounter or on resident microglia for Fc mediated uptake. Further, while microglia may become dysfunctional with age (Mosher and Wyss-Coray, 2014), CAR-Ms could theoretically be engineered from allogeneic young healthy donors and overcome some of the functional challenges facing aged cells.

The ability of CAR-Ms to degrade amyloid may also lead to reduced incidence of amyloid-related imaging abnormalities (ARIA) compared to antibody therapy. ARIA is thought to occur in the setting of anti-Aβ monoclonal antibody therapy when amyloid-antibody complexes are cleared via perivascular clearance mechanisms and accumulate in perivascular spaces, which can impair further perivascular drainage of these complexes and cause inflammatory reactions that damage the brain vasculature and disrupt the blood-brain-barrier, resulting in edema and microhemorrhage (Greenberg et al., 2020; Sperling et al., 2012). Future studies will determine if CAR-Ms can reduce amyloid plaque burden with less incidence of ARIA than anti-Aβ monoclonal antibody therapy.

There are several important limitations to these studies. For poorly understood reasons, CAR-Ms injected into the hippocampus of APP/PS1 mice were limited in their survival and migration within the brain niche. We have not examined why the CAR-Ms fail to persist long-term within the brain, but several possibilities exist. Just as microglia have different phenotypes throughout the brain due to differences in local microenvironment (Spiteri et al., 2022), fully differentiated CAR-Ms may lack the complete set of receptors necessary to thrive within the brain niche. Even if they can survive within the brain, they may become out-competed by more fit microglia as they recover from preconditioning depletion. Alternatively, the CAR-Ms may be immunologically depleted, due to either MHC expression of foreign antigenic peptides (such as from GFP-Luciferase), or due to mixed strain immunoreactivity (APP/PS1 mice are on a mixed C57BL/6 × C3H background, since pure C57BL/6 APP/PS1 mice develop seizures at a considerable rate) (Minkeviciene et al., 2009). Additional studies are needed to determine factors that promote the migration of peripheral myeloid cells within the brain parenchyma. While the mechanism of failure of long-term CAR-M engraftment into the brain is of interest academically, it is not clear whether achieving long-term engraftment would be desired clinically. The fact that these CAR-Ms are allogeneic and do not persist long-term, but still have a significant local effect *in vivo* is encouraging for the possibility that if CAR-Ms were to become therapeutically relevant for AD in the future, an allogeneic product may be feasible. Another limitation of this study is that CAR-Ms were delivered intrahippocampally. Clinical translation of Aβ CAR-Ms will likely require that these cells be delivered intravenously, but in the context of AD, little is known about the factors that promote peripheral monocyte/macrophage entry into the brain. Our previous work showed that peripheral monocytes can enter the AD brain in APP/PS1 mice and reduce plaque load in the absence of any intervention (Yan et al., 2022), suggesting that increasing the number of peripheral monocytes entering the brain, especially Aβ CAR-modified ones, will further reduce plaque load. Identification of factors driving peripheral monocyte infiltration into the AD brain requires additional study but will inform further rounds of CAR engineering to promote greater infiltration into the AD brain and enhance the translatability and efficacy of this therapy. Another potential strategy for clinical translation is the delivery of anti-Aβ CAR mRNA with monocyte/macrophage-targeted lipid nanoparticles, as has been done for CAR-T cells in the setting of cardiac fibrosis (Rurik et al., 2022).

Although we use Aβ as the target of these CAR-Ms for proof of principle, theoretically any other protein may be targeted and degraded by simply replacing the scFv domain of the CAR. Our data establish CAR-Ms as one additional potential approach in the therapeutic toolbox for AD, and extend the potential use of CAR-Ms beyond cancer.

## Materials and Methods

### Plasmids

The Aβ CAR construct was generated by adding the aducanumab scFv, and mFcγR extracellular, transmembrane, and intracellular domains to a MuLV retroviral backbone. The control CAR construct was generated by adding an anti-EphA2 scFv and CD8 extracellular transmembrane domains to the retroviral backbone. The GFP-luc construct was made from GFP and F-luciferase in a MuLV retroviral backbone. The M-CSF construct was generated by adding Thy1.1 and M-CSF to the retroviral backbone.

### Cell lines

RD114 and Plat-E cells were cultured in DMEM media (Sigma-Aldrich) with 10% FBS (Atlas Biologicals), 1% penicillin and streptomycin (Gibco). The generation of HoxB8 cells has been previously described (Redecke et al., 2013). Briefly, femoral bone marrow cells from C57BL/6J mice were isolated and cultured in recombinant mouse IL-3, IL-6, and SCF for 2 days. On the third day of culture, cells were cultured in media with FLT3-ligand and infected with retroviral introduction of an MSCV vector containing the mouse Hoxb8 DNA and a human estrogen receptor binding domain with a mutation that prevents physiological concentrations of estrogen from binding. HoxB8 cells were maintained in RPMI media (Gibco) with 10% FBS (Atlas Biologicals), 1% penicillin and streptomycin (Gibco), 1%sodium pyruvate (Gibco), non-essential amino acids (Gibco), HEPES (Thermo-Fischer), 2Mm L-glutamine, 0.1% beta-mercaptoethanol (Gibco), FLT3-ligand and beta estradiol (Sigma). All the HoxB8 cells were transduced with retroviral vectors obtained from Plat-E cell lines. After transduction, FACS sorting is performed to obtain 100% retroviral vector+ HoxB8 cells. To induce differentiation, sorted HoxB8 cells were incubated in RPMI media supplemented with 10% M-CSF for 6 days. Macrophages were lifted from cell culture plates for downstream assays with Accutase (Innovative Cell Technologies). All the cell lines were regularly tested for mycoplasma by in-house PCR.

### Cell transfection and transduction

The RD114 retroviral packaging cell line was seeded to 6-well plates at a concentration of 500,000 cells/well the day before transfection. 2.5 µg of plasmid DNA was mixed with 5 µL Polyethyleneimine (PEI) at 1ug/ml stock solution (Alfa Aesar) and incubated for 15 min at room temperature before being added to the RD114 cells. 16-24 hours later, media was changed and retrovirus was collected for use starting 24 hours later. RD114 virus was used to generate stable virus-producer Plat-E cell lines generated via retroviral transduction with 8ug/ml polybrene and spinfection at 3000rpm for 1 hr. Plat-E virus was used to transduce HoxB8 cells, which were transduced with 4ug/ml polybrene and spinfection at 600xg for 30mins.

### Flow cytometry

All antibodies were titrated. CAR expression was measured with Goat anti-human IgG Alexa Fluor 647 (156339, Jackson ImmunoResearch) and transduction with the M-CSF construct was measured with anti-Thy1.1 PE (B336285, BioLegend). M-CSF differentiated macrophages were stained with Fc block and Oligoblock (made in-house), anti-F4/80 BV421 (0351659, BD BioScience) and anti-CD64 BV786 (0328308, BD BioScience). Live/dead staining and the M-CSF CAR-M viability assay was conducted with ZombieNIR Fixable Viabilty Dye (423106, Biolegend). Flow cytometry data was acquired on a Cytek NL-3000, and data was analyzed with FlowJo version 10.8.1 (Treestar/BD Biosciences).

### *In vitro* phagocytosis assay

CAR-macrophages were generated by differentiating HoxB8 cells with M-CSF for 6 days and were then seeded in 96-well plates at a concentration of 20,000/well in 200 µL serum free RPMI. 1 µL of HiLyte™ Fluor 488-labeled Amyloid-beta (1-42) (Anaspec) prepared by reconstituting 0.1mg in 50 µL of 1% NH_4_OH, was added into each well. After 2 or 4 hours of co-incubation, the wells were washed with PBS and analyzed by flow cytometry.

### *Ex vivo* phagocytosis assay

12-14 month old APP/PS1 mice were perfused with PBS prior to brain extraction. Brains were frozen on dry ice and then stored at −80°C. Prior to each assay, frozen brains were sectioned into 10 µm slices using a CryoStat, and subsequently placed on PDL coated coverslips. The coverslips were then placed in cell culture plates immersed in culture media.

CAR Hoxb8 cells were induced to differentiate to macrophages in RPMI media with M-CSF for 6 days. On day 6, floating cells were washed away with PBS and adherent CAR-Macrophages were lifted with Accutase (Innovative Cell Technologies). Brain slices were incubated with 2×10^5^ control or Aβ CAR-Ms in 1mL of differentiation media (RPMI + M-CSF) for 44 hours. Cells were cultured on adjacent brain slices to approximately match plaque load and distribution. After incubation, brain slices were fixed in 4% PFA for 20 min. Slices were then stained with X-34 dye or HJ3.4 antibody. High resolution images of the slices were taken using a NanoZoomer Digital Scanner (Hamamatsu Photonics). The total area of plaque coverage was measured with NIH ImageJ software and expressed as percentage total area. The frequency of size classified plaques was analyzed with ImageJ software.

### Intracranial injection and BLI imaging

APP/PS1 mice (B6;C3-Tg(APPswe,PSEN1dE9)85Dbo/Mmjax from Jackson Labs, MMRRC Stock No. 34928, maintained as C57BL/6 x C3H strain), 12-14 month of age, were preconditioned with either 20mg/kg Busulfan once a day for 2 days or PLX5622 (HY-114153, MedChemExpress) at 50mg/kg twice daily for 4 days by intraperitoneal (IP) injection. Busulfan was dissolved in DMSO (Sigma) at a concentration of 30 mg/mL and diluted to 3mg/mL in PBS for injection. PLX5622 was dissolved in DMSO at a concentration of 50mg/mL and diluted to 5mg/mL in in 20% Kolliphor RH40 (Sigma-Aldrich) in PBS.

Cells were prepared for intracranial injection by incubating CAR HoxB8 cells in 10% M-CSF differentiation media for 6 days. On Day 6 post differentiation, the macrophages were lifted with Accutase, washed 3 times with PBS, and loaded into a 5 µL Hamilton syringe at a concentration of 1.5×10^5^ cells/µL. 2µL of Aβ CAR-Ms were injected into the right hippocampus and 2µL of PBS or control CAR-Ms were injected into the left hippocampus (Coordinate: AP: −2.0, ML:

±1.6 DV: −1.5). CAR-Ms were injected with 200 ng/ml recombinant M-CSF. Bioluminescence imaging (BLI) was performed every 3-7 days; mouse heads were shaved before imaging. Mice were grouped in three day intervals when charting BLI over time.

### Immunohistochemistry for *in vivo* studies

12-14 month old male and female APP/PS1 mice were sacrificed on day 14 with Fatal-Plus and perfused with PBS. For microglia depletion studies (Fig. S3c), mice were sacrificed on day 4 post preconditioning with PBS, Busulfan, or PLX5622 as above. Brains were removed, fixed for 24 hours in 4% paraformaldehyde fixative in 0.1 M phosphate buffer (PB) (pH 7.4) and then transferred to a solution containing 30% sucrose in 0.1 M PB until the tissue sank down. The brain was then sectioned (40 µm) and twelve equally spaced sections (80 µm apart) containing the dorsal part of hippocampus were immunostained using antibodies against GFP and HJ3.4 delineated below. For microglia depletion studies, brain sections were stained with Iba1. Brain sections were permeabilized and blocked with 0.3% Triton X-100/ 3% dry milk in 0.01 M PBS for 30 minutes followed by incubation with primary antibodies overnight at 4°C and fluorescently labeled secondary antibodies at 37°C for 1 hour. Primary and secondary antibodies employed are shown in Supplemental Table 1. For X-34 staining, brain slices were mounted on glass slides. Tissue was permeabilized with 0.25% Triton for 30 minutes and stained with X-34 dissolved in a solution of 40% ethanol in water, pH 10, for 20 minutes. The tissue was then rinsed in distilled water and mounted.

### Assessment of Microglial Depletion with Preconditioning

Images were acquired with a Biotek Cytation 5 (Agilent) and images were analyzed with Biotek Gen5 software. For each mouse, six regions of interest (ROIs) were captured in each hippocampus at 20x magnification, for a total of 12 ROIs per mouse and three mice per group. In each ROI, Iba-1 positive cells were counted using the following parameters in Gen5: threshold minimum 4800, minimum size 10 µm, maximum size 60 µm, include primary edge objects, do not split touching objects. This mask was applied using the same settings to all ROIs across conditions to quantify microglia in the hippocampus.

### Assessment of amyloid plaques

Images were acquired using a Zeiss Axio Scan Z1 or a Nikon AXR Confocal Microscope. For each mouse, five to nine sections containing the dorsal hippocampus (spaced 80 µm apart) were analyzed using Image J. Circular regions of interest (ROIs) with uniform areas of 3622 um^2^ were created, centered around the injection site in the hippocampus (residual GFP signal). Plaque load was quantified within these circles using brain sections stained with either HJ3.4 or X-34. Plaque load was expressed as percentage of area within the ROI. Uniform circle sizes were used across all slices. For controls, circular ROIs in mirror cortical regions were analyzed to ensure the density of plaques was relatively uniform across mirror regions of each hemisphere.

### Statistics

Statistical analyses were performed in GraphPad Prism version 9.5.1. Results are expressed as mean ± s.e.m. Statistical differences were assessed with the unpaired 2-tailed Student’s t test for comparison of 2 experimental groups or ANOVA for 3+ experimental groups. P values from ANOVA with multiple-comparisons were generated with Tukey’s multiple-comparisons test. Specific test used in each case is denoted in figure legends. Unless otherwise stated, *P < 0.05, **P < 0.01, ***P < 0.001, ****P < 0.0001. A 2-tailed P value of less than 0.05 was considered statistically significant.

### Study approval

All animal studies were approved by the IACUC at Washington University School of Medicine.

## Acknowledgements

The authors would like to thank E. Gonzales and S. Grathwohl of the Hope Center Animal Surgery Core for their help with intrahippocampal injections and animal care as well as the Washington University School of Medicine’s Department of Pathology and Immunology’s Flow Cytometry & Fluorescence Activated Cell Sorting Core for their assistance with cell sorting. Figures 1a, 2a, 3a, 3c, 4, and s2a were created using BioRender.

This work was supported by grants from the NIH (R01 NS094692 to JML and AD, R21 NS082529 to JML and AD, R01 AG079503 to JML, and DP5OD026427 to CJD).

---

**Supplemental Figure 1:**
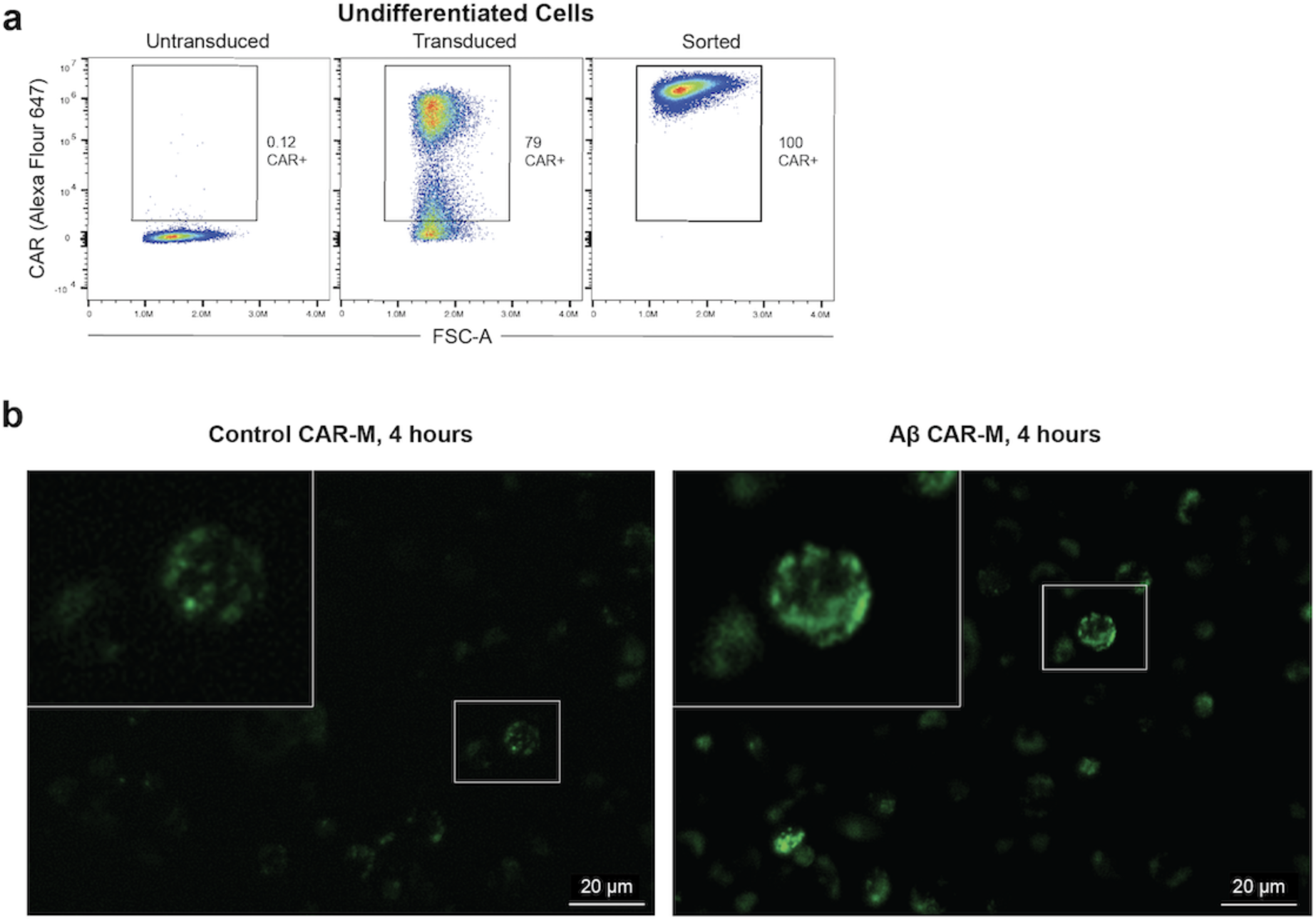
Generation and validation of CAR HoxB8 cells. **a)** Representative FACS plots showing surface expression of the Aβ CAR on untransduced, retrovirally transduced, and transduced and sorted HoxB8 cells used for downstream experiments. Numbers represent percentage of cells in the indicated gate. Representative of n>3 independent experiments. **b)** Representative images of control or Aβ CAR Ms co-cultured with AF488 tagged Aβ(1-42) for 4 hours.

**Supplemental Figure 2:**
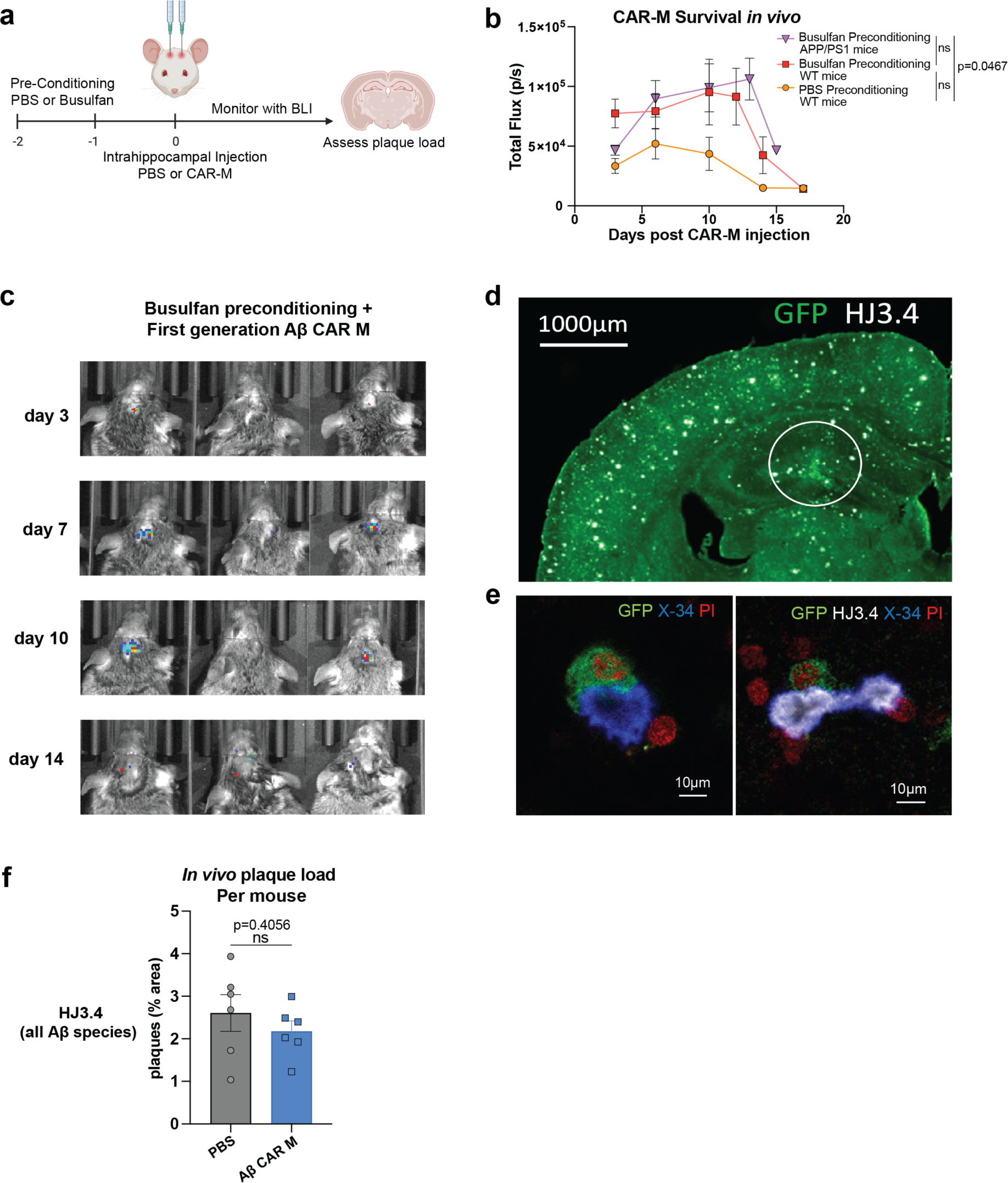
First generation Aβ CAR-Ms have limited expansion and survival and fail to reduce plaque load *in vivo* when administered with Busulfan preconditioning. **a)** Schematic of Busulfan pre-conditioning and intrahippocampal injection of Aβ CAR-Ms. **b)** Non-invasive bioluminescence imaging tracking CAR-M persistence after intrahippocampal injection. n=6-14 mice per group. Statistical significance calculated with one-way ANOVA with Tukey’s multiple comparisons test. **c)** Representative bioluminescence images following Busulfan preconditioning and intrahippocampal injection of Aβ CAR-Ms. Days indicates days post-intrahippocampal injection. **d)** Representative immunofluorescence microscopy image showing GFP+ CAR-Ms localized to the hippocampus 13 days after intrahippocampal injection. Circular region of interest indicates area in which plaque load was quantified. **e)** Representative immunofluorescence microscopy images of Aβ CAR-Ms binding to amyloid plaque *in vivo*. **f)** Assessment of plaque load after intrahippocampal injection of PBS or Aβ CAR-M in n=6 aged APP/PS1 mice. Mice were sacrificed on day 14 post intrahippocampal injection and brain tissue was sectioned and stained with HJ3.4 to assess plaque load. Data shown as mean ± s.e.m. Statistical significance was calculated with unpaired t-tests.

**Supplemental Figure 3:**
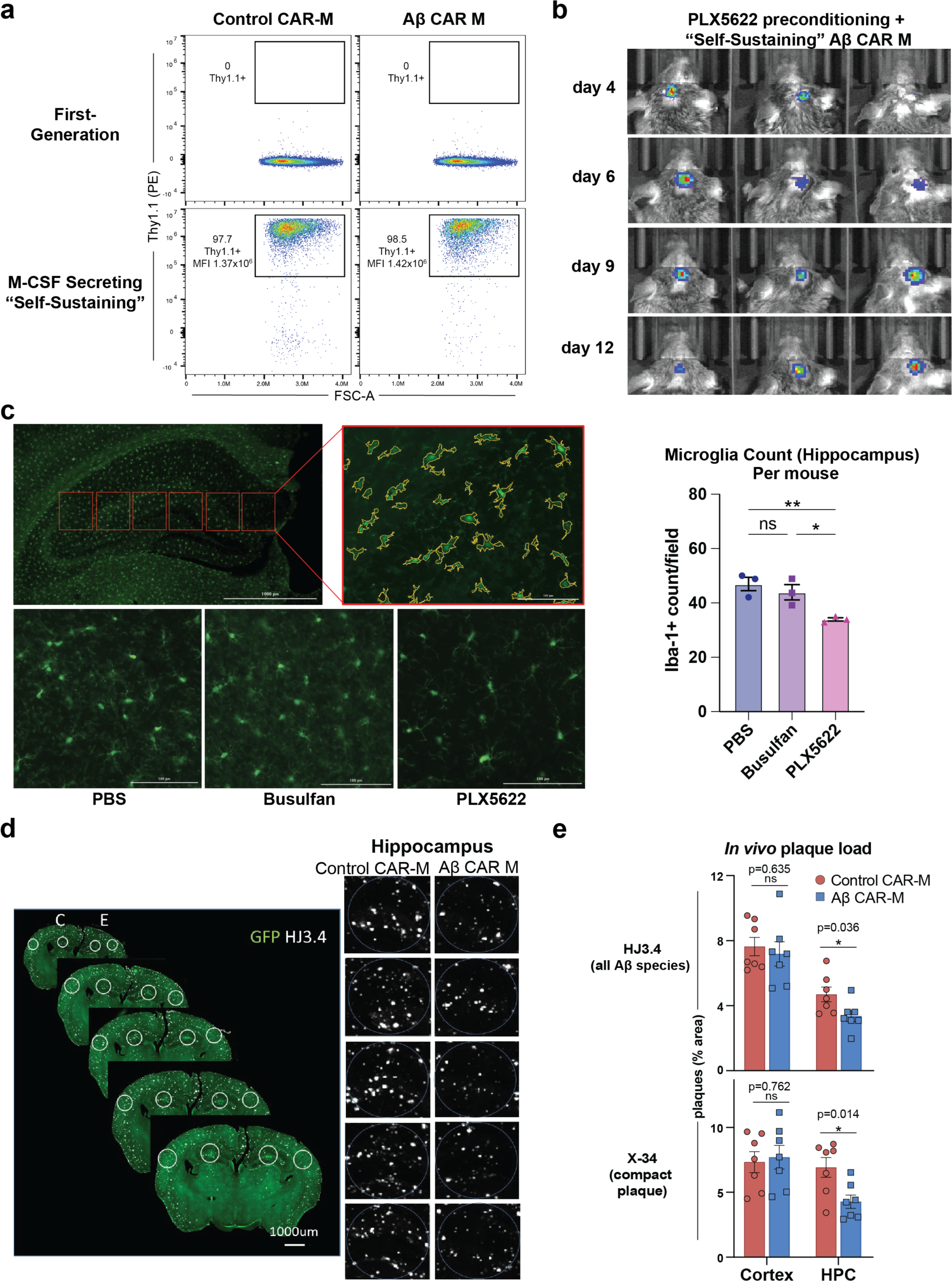
Self-sustaining, M-CSF secreting Aβ CAR-Ms expand *in vivo* when administered with PLX5622 preconditioning and reduce plaque load in the locally in the hippocampus *in vivo*. **a)** Representative FACS plots showing surface expression of Thy1.1 on control and Aβ CAR-Ms before and after retroviral transduction of HoxB8 cells with the M-CSF construct and sorting for Thy1.1+ cells. Gated on single, live, CAR+ cells. MFI, mean fluorescence intensity **b)** Representative bioluminescence images following PLX5622 preconditioning and intrahippocampal injection of self-sustaining Aβ CAR-Ms. Days indicates days post-intrahippocampal injection. **c)** (left upper) Representative image showing the six regions of interest (ROI) in which Iba-1 positive cells were quantified in each hippocampus and a higher power view of one ROI showing the cell masking used for quantification. (left lower) Representative images showing one ROI from mice treated with PBS for 2 days, 20 mg/kg Busulfan for 2 days, or 50 mg/kg PLX5622 for 4 days, twice a day. (right) quantification of Iba-1+ cells where each dot represents the average Iba-1+ cell count in all ROIs per mouse. Statistical significance was calculated with a one-way ANOVA with Tukey’s multiple comparisons test. **d)** Representative images of brain sections from CAR-M treated aged APP/PS1 mice stained with HJ3.4. Images indicate circular regions of interest centered around GFP signal in the hippocampus representing the cell injection area in which plaque was quantified, highlighted in higher magnification on the right. Control regions of interest were quantified in the cortex to ensure uniform plaque load between mice. “C”= control CAR-M treated side, “E”= Aβ CAR-M treated side. **e)** Assessment of plaque load in the cortex and the hippocampus (HPC) after intrahippocampal injection of self-sustaining control or Aβ CAR-Ms in n=7 aged APP/PS1 mice. Mice were sacrificed on day 12 or 13 post intrahippocampal injection and brain tissue was sectioned and stained with HJ3.4 or X-34 to assess plaque load in the regions of interest shown in **d)**. Data shown as mean ± s.e.m. Statistical significance was calculated with unpaired t-tests. *P < 0.05, **P < 0.01, ***P < 0.001, ****P < 0.0001, ns, not significant.

**Supplemental Table 1:**

| Antigen | Primary working concentration | Source | Secondary working concentration | Source |
| --- | --- | --- | --- | --- |
| Iba1 | 1:1000 | Rabbit anti-Iba1 (Wako Pure Chemical Industries, Ltd. cat# 019-19741) | 1:800 | Anti-rabbit Alexa488 (Thermo Fisher) |
| GFP | 1:500 | Goat anti-GFP (Rockland antibodies) | 1:800 | Anti-goat Alexa488 (Life Sciences) |
| Aβ (HJ3.4) | 1:1000 | Biotinylated antibody, gift from Dr. David M. Holtzman | 1:800 | Cy5 Streptavidin (Jackson ImmunoResearch Inc.) |

## References

1. Ajami, B., J.L. Bennett, C. Krieger, W. Tetzlaff, and F.M. Rossi. 2007. Local self-renewal can sustain CNS microglia maintenance and function throughout adult life. Nat Neurosci 10:1538–1543.

2. Bacskai, B.J., S.T. Kajdasz, R.H. Christie, C. Carter, D. Games, P. Seubert, D. Schenk, and B.T. Hyman. 2001. Imaging of amyloid-beta deposits in brains of living mice permits direct observation of clearance of plaques with immunotherapy. Nat Med 7:369–372.

3. Bard, F., C. Cannon, R. Barbour, R.L. Burke, D. Games, H. Grajeda, T. Guido, K. Hu, J. Huang, K. Johnson-Wood, K. Khan, D. Kholodenko, M. Lee, I. Lieberburg, R. Motter, M. Nguyen, F. Soriano, N. Vasquez, K. Weiss, B. Welch, P. Seubert, D. Schenk, and T. Yednock. 2000. Peripherally administered antibodies against amyloid beta-peptide enter the central nervous system and reduce pathology in a mouse model of Alzheimer disease. Nat Med 6:916-919.

4. Basilico, B., L. Ferrucci, P. Ratano, M.T. Golia, A. Grimaldi, M. Rosito, V. Ferretti, I. Reverte, C. Sanchini, M.C. Marrone, M. Giubettini,V. De Turris, D. Salerno, S. Garofalo, M.K. St-Pierre, M. Carrier, M. Renzi, F. Pagani, B. Modi, M. Raspa, F. Scavizzi, C.T. Gross, S. Marinelli, M.E. Tremblay, D. Caprioli, L. Maggi, C. Limatola, S. Di Angelantonio, and D. Ragozzino. 2022. Microglia control glutamatergic synapses in the adult mouse hippocampus. Glia 70:173–195.

5. Boissonneault, V., M. Filali, M. Lessard, J. Relton, G. Wong, and S. Rivest. 2009. Powerful beneficial effects of macrophage colony-stimulating factor on beta-amyloid deposition and cognitive impairment in Alzheimer’s disease. Brain 132:1078–1092.

6. Budd Haeberlein, S., P.S. Aisen, F. Barkhof, S. Chalkias, T. Chen, S. Cohen, G. Dent, O. Hansson, K. Harrison, C. von Hehn, T. Iwatsubo, C. Mallinckrodt, C.J. Mummery, K.K. Muralidharan, I. Nestorov, L. Nisenbaum, R. Rajagovindan, L. Skordos, Y. Tian, C.H. van Dyck, B. Vellas, S. Wu, Y. Zhu, and A. Sandrock. 2022. Two Randomized Phase 3 Studies of Aducanumab in Early Alzheimer’s Disease. J Prev Alzheimers Dis 9:197–210.

7. Champion, J.A., A. Walker, and S. Mitragotri. 2008. Role of particle size in phagocytosis of polymeric microspheres. Pharm Res 25:1815–1821.

8. Couzin-Frankel, J., and C. Piller. 2022. Alzheimer’s drug stirs excitement-and concerns. Science 378:1030–1031.

9. d’Errico, P., S. Ziegler-Waldkirch, V. Aires, P. Hoffmann, C. Mezo, D. Erny, L.S. Monasor, S. Liebscher, V.M. Ravi, K. Joseph, O. Schnell, K. Kierdorf, O. Staszewski, S. Tahirovic, M. Prinz, and M. Meyer-Luehmann. 2022. Microglia contribute to the propagation of Abeta into unaffected brain tissue. Nat Neurosci 25:20–25.

10. DeMattos, R.B., K.R. Bales, D.J. Cummins, J.C. Dodart, S.M. Paul, and D.M. Holtzman. 2001. Peripheral anti-A beta antibody alters CNS and plasma A beta clearance and decreases brain A beta burden in a mouse model of Alzheimer’s disease. Proc Natl Acad Sci U S A 98:8850–8855.

11. Drieu, A., S. Du, S.E. Storck, J. Rustenhoven, Z. Papadopoulos, T. Dykstra, F. Zhong, K. Kim, S. Blackburn, T. Mamuladze, O. Harari, C.M. Karch, R.J. Bateman, R. Perrin, M. Farlow, J. Chhatwal, N. Dominantly Inherited Alzheimer, S. Hu, G.J. Randolph, I. Smirnov, and J. Kipnis. 2022. Parenchymal border macrophages regulate the flow dynamics of the cerebrospinal fluid. Nature 611:585–593.

12. Fiala, M., J. Lin, J. Ringman, V. Kermani-Arab, G. Tsao, A. Patel, A.S. Lossinsky, M.C. Graves, A. Gustavson, J. Sayre, E. Sofroni, T. Suarez, F. Chiappelli, and G. Bernard. 2005. Ineffective phagocytosis of amyloid-beta by macrophages of Alzheimer’s disease patients. J Alzheimers Dis 7:221–232; discussion 255-262.

13. Frenkel, D., M. Balass, E. Katchalski-Katzir, and B. Solomon. 1999. High affinity binding of monoclonal antibodies to the sequential epitope EFRH of beta-amyloid peptide is essential for modulation of fibrillar aggregation. J Neuroimmunol 95:136–142.

14. Funk, K.E., H. Mirbaha, H. Jiang, D.M. Holtzman, and M.I. Diamond. 2015. Distinct Therapeutic Mechanisms of Tau Antibodies: Promoting Microglial Clearance Versus Blocking Neuronal Uptake. J Biol Chem 290:21652–21662.

15. Gattinoni, L., S.E. Finkelstein, C.A. Klebanoff, P.A. Antony, D.C. Palmer, P.J. Spiess, L.N. Hwang, Z. Yu, C. Wrzesinski, D.M. Heimann, C.D. Surh, S.A. Rosenberg, and N.P. Restifo. 2005. Removal of homeostatic cytokine sinks by lymphodepletion enhances the efficacy of adoptively transferred tumor-specific CD8+ T cells. J Exp Med 202:907–912.

16. Giuffrida, L., K. Sek, M.A. Henderson, I.G. House, J. Lai, A.X.Y. Chen, K.L. Todd, E.V. Petley, S. Mardiana, I. Todorovski, E. Gruber, M.J. Kelly, B.J. Solomon, S.J. Vervoort, R.W. Johnstone, I.A. Parish, P.J. Neeson, L.M. Kats, P.K. Darcy, and P.A. Beavis. 2020. IL-15 Preconditioning Augments CAR T Cell Responses to Checkpoint Blockade for Improved Treatment of Solid Tumors. Mol Ther 28:2379–2393.

17. Grathwohl, S.A., R.E. Kalin, T. Bolmont, S. Prokop, G. Winkelmann, S.A. Kaeser, J. Odenthal, R. Radde, T. Eldh, S. Gandy, A. Aguzzi, M. Staufenbiel, P.M. Mathews, H. Wolburg, F.L. Heppner, and M. Jucker. 2009. Formation and maintenance of Alzheimer’s disease beta-amyloid plaques in the absence of microglia. Nat Neurosci 12:1361–1363.

18. Greenberg, S.M., B.J. Bacskai, M. Hernandez-Guillamon, J. Pruzin, R. Sperling, and S.J. van Veluw. 2020. Cerebral amyloid angiopathy and Alzheimer disease - one peptide, two pathways. Nat Rev Neurol 16:30–42.

19. Henry, R.J., R.M. Ritzel, J.P. Barrett, S.J. Doran, Y. Jiao, J.B. Leach, G.L. Szeto, J. Wu, B.A. Stoica, A.I. Faden, and D.J. Loane. 2020. Microglial Depletion with CSF1R Inhibitor During Chronic Phase of Experimental Traumatic Brain Injury Reduces Neurodegeneration and Neurological Deficits. J Neurosci 40:2960–2974.

20. Hickman, S.E., E.K. Allison, and J. El Khoury. 2008. Microglial dysfunction and defective beta-amyloid clearance pathways in aging Alzheimer’s disease mice. J Neurosci 28:8354–8360.

21. Huang, Y., K.E. Happonen, P.G. Burrola, C. O’Connor, N. Hah, L. Huang, A. Nimmerjahn, and G. Lemke. 2021. Microglia use TAM receptors to detect and engulf amyloid beta plaques. Nat Immunol 22:586–594.

22. Karran, E., and B. De Strooper. 2022. The amyloid hypothesis in Alzheimer disease: new insights from new therapeutics. Nat Rev Drug Discov 21:306–318.

23. Klichinsky, M., M. Ruella, O. Shestova, X.M. Lu, A. Best, M. Zeeman, M. Schmierer, K. Gabrusiewicz, N.R. Anderson, N.E. Petty, K.D. Cummins, F. Shen, X. Shan, K. Veliz, K. Blouch, Y. Yashiro-Ohtani, S.S. Kenderian, M.Y. Kim, R.S. O’Connor, S.R. Wallace, M.S. Kozlowski, D.M. Marchione, M. Shestov, B.A. Garcia, C.H. June, and S. Gill. 2020. Human chimeric antigen receptor macrophages for cancer immunotherapy. Nat Biotechnol 38:947–953.

24. Klyubin, I., D.M. Walsh, C.A. Lemere, W.K. Cullen, G.M. Shankar, V. Betts, E.T. Spooner, L. Jiang, R. Anwyl, D.J. Selkoe, and M.J. Rowan. 2005. Amyloid beta protein immunotherapy neutralizes Abeta oligomers that disrupt synaptic plasticity in vivo. Nat Med 11:556–561.

25. Lamb, Y.N. 2019. Pexidartinib: First Approval. Drugs 79:1805–1812.

26. Lucin, K.M., C.E. O’Brien, G. Bieri, E. Czirr, K.I. Mosher, R.J. Abbey, D.F. Mastroeni, J. Rogers, B. Spencer, E. Masliah, and T. Wyss-Coray. 2013. Microglial beclin 1 regulates retromer trafficking and phagocytosis and is impaired in Alzheimer’s disease. Neuron 79:873–886.

27. Mailankody, S., S.M. Devlin, J. Landa, K. Nath, C. Diamonte, E.J. Carstens, D. Russo, R. Auclair, L. Fitzgerald, B. Cadzin, X. Wang, D. Sikder, B. Senechal, V.P. Bermudez, T.J. Purdon, K. Hosszu, D.P. McAvoy, T. Farzana, E. Mead, J.A. Wilcox, B.D. Santomasso, G.L. Shah, U.A. Shah, N. Korde, A. Lesokhin, C.R. Tan, M. Hultcrantz, H. Hassoun, M. Roshal, F. Sen, A. Dogan, O. Landgren, S.A. Giralt, J.H. Park, S.Z. Usmani, I. Riviere, R.J. Brentjens, and E.L. Smith. 2022. GPRC5D-Targeted CAR T Cells for Myeloma. N Engl J Med 387:1196–1206.

28. Marchetti, L., and B. Engelhardt. 2020. Immune cell trafficking across the blood-brain barrier in the absence and presence of neuroinflammation. Vasc Biol 2:H1-H18.

29. Minkeviciene, R., S. Rheims, M.B. Dobszay, M. Zilberter, J. Hartikainen, L. Fulop, B. Penke, Y. Zilberter, T. Harkany, A. Pitkanen, and H. Tanila. 2009. Amyloid beta-induced neuronal hyperexcitability triggers progressive epilepsy. J Neurosci 29:3453–3462.

30. Mosher, K.I., and T. Wyss-Coray. 2014. Microglial dysfunction in brain aging and Alzheimer’s disease. Biochem Pharmacol 88:594–604.

31. Murad, J.P., D. Tilakawardane, A.K. Park, L.S. Lopez, C.A. Young, J. Gibson, Y. Yamaguchi, H.J. Lee, K.T. Kennewick, B.J. Gittins, W.C. Chang, C.P. Tran, C. Martinez, A.M. Wu, R.E. Reiter, T.B. Dorff, S.J. Forman, and S.J. Priceman. 2021. Pre-conditioning modifies the TME to enhance solid tumor CAR T cell efficacy and endogenous protective immunity. Mol Ther 29:2335–2349.

32. Neelapu, S.S., F.L. Locke, N.L. Bartlett, L.J. Lekakis, D.B. Miklos, C.A. Jacobson, I. Braunschweig, O.O. Oluwole, T. Siddiqi, Y. Lin, J.M. Timmerman, P.J. Stiff, J.W. Friedberg, I.W. Flinn, A. Goy, B.T. Hill, M.R. Smith, A. Deol, U. Farooq, P. McSweeney, J. Munoz, I. Avivi, J.E. Castro, J.R. Westin, J.C. Chavez, A. Ghobadi, K.V. Komanduri, R. Levy, E.D. Jacobsen, T.E. Witzig, P. Reagan, A. Bot, J. Rossi, L. Navale, Y. Jiang, J. Aycock, M. Elias, D. Chang, J. Wiezorek, and W.Y. Go. 2017. Axicabtagene Ciloleucel CAR T-Cell Therapy in Refractory Large B-Cell Lymphoma. N Engl J Med

33. Pacheco, P., D. White, and T. Sulchek. 2013. Effects of microparticle size and Fc density on macrophage phagocytosis. PLoS One 8:e60989.

34. Park, J.H., I. Riviere, M. Gonen, X. Wang, B. Senechal, K.J. Curran, C. Sauter, Y. Wang, B. Santomasso, E. Mead, M. Roshal, P. Maslak, M. Davila, R.J. Brentjens, and M. Sadelain. 2018. Long-Term Follow-up of CD19 CAR Therapy in Acute Lymphoblastic Leukemia. N Engl J Med 378:449–459.

35. Pratten, M.K., and J.B. Lloyd. 1986. Pinocytosis and phagocytosis: the effect of size of a particulate substrate on its mode of capture by rat peritoneal macrophages cultured in vitro. Biochim Biophys Acta 881:307–313.

36. Raje, N., J. Berdeja, Y. Lin, D. Siegel, S. Jagannath, D. Madduri, M. Liedtke, J. Rosenblatt, M.V. Maus, A. Turka, L.P. Lam, R.A. Morgan, K. Friedman, M. Massaro, J. Wang, G. Russotti, Z. Yang, T. Campbell, K. Hege, F. Petrocca, M.T. Quigley, N. Munshi, and J.N. Kochenderfer. 2019. Anti-BCMA CAR T-Cell Therapy bb2121 in Relapsed or Refractory Multiple Myeloma. N Engl J Med 380:1726–1737.

37. Redecke, V., R. Wu, J. Zhou, D. Finkelstein, V. Chaturvedi, A.A. High, and H. Hacker. 2013. Hematopoietic progenitor cell lines with myeloid and lymphoid potential. Nat Methods 10:795–803.

38. Reish, N.J., P. Jamshidi, B. Stamm, M.E. Flanagan, E. Sugg, M. Tang, K.L. Donohue, M. McCord, C. Krumpelman, M.M. Mesulam, R. Castellani, and S.H. Chou. 2023. Multiple Cerebral Hemorrhages in a Patient Receiving Lecanemab and Treated with t-PA for Stroke. N Engl J Med 388:478–479.

39. Rurik, J.G., I. Tombacz, A. Yadegari, P.O. Mendez Fernandez, S.V. Shewale, L. Li, T. Kimura, O.Y. Soliman, T.E. Papp, Y.K. Tam, B.L. Mui, S.M. Albelda, E. Pure, C.H. June, H. Aghajanian, D. Weissman, H. Parhiz, and J.A. Epstein. 2022. CAR T cells produced in vivo to treat cardiac injury. Science 375:91–96.

40. Sailor, K.A., G. Agoranos, S. Lopez-Manzaneda, S. Tada, B. Gillet-Legrand, C. Guerinot, J.B. Masson, C.L. Vestergaard, M. Bonner, K. Gagnidze, G. Veres, P.M. Lledo, and N. Cartier. 2022. Hematopoietic stem cell transplantation chemotherapy causes microglia senescence and peripheral macrophage engraftment in the brain. Nat Med 28:517–527.

41. Salloway, S., S. Chalkias, F. Barkhof, P. Burkett, J. Barakos, D. Purcell, J. Suhy, F. Forrestal, Y. Tian, K. Umans, G. Wang, P. Singhal, S. Budd Haeberlein, and K. Smirnakis. 2022. Amyloid-Related Imaging Abnormalities in 2 Phase 3 Studies Evaluating Aducanumab in Patients With Early Alzheimer Disease. JAMA Neurol 79:13-21.

42. Salter, M.W., and B. Stevens. 2017. Microglia emerge as central players in brain disease. Nat Med 23:1018–1027.

43. Scheltens, P., B. De Strooper, M. Kivipelto, H. Holstege, G. Chetelat, C.E. Teunissen, J. Cummings, and W.M. van der Flier. 2021. Alzheimer’s disease. Lancet 397:1577–1590.

44. Schenk, D., R. Barbour, W. Dunn, G. Gordon, H. Grajeda, T. Guido, K. Hu, J. Huang, K. Johnson-Wood, K. Khan, D. Kholodenko, M. Lee, Z. Liao, I. Lieberburg, R. Motter, L. Mutter, F. Soriano, G. Shopp, N. Vasquez, C. Vandevert, S. Walker, M. Wogulis, T. Yednock, D. Games, and P. Seubert. 1999. Immunization with amyloid-beta attenuates Alzheimer-disease-like pathology in the PDAPP mouse. Nature 400:173–177.

45. Sevigny, J., P. Chiao, T. Bussiere, P.H. Weinreb, L. Williams, M. Maier, R. Dunstan, S. Salloway, T. Chen, Y. Ling, J. O’Gorman, F. Qian, M. Arastu, M. Li, S. Chollate, M.S. Brennan, O. Quintero-Monzon, R.H. Scannevin, H.M. Arnold, T. Engber, K. Rhodes, J. Ferrero, Y. Hang, A. Mikulskis, J. Grimm, C. Hock, R.M. Nitsch, and A. Sandrock. 2016. The antibody aducanumab reduces Abeta plaques in Alzheimer’s disease. Nature 537:50–56.

46. Shibuya, Y., K.K. Kumar, M.M. Mader, Y. Yoo, L.A. Ayala, M. Zhou, M.A. Mohr, G. Neumayer, I. Kumar, R. Yamamoto, P. Marcoux, B. Liou, F.C. Bennett, H. Nakauchi, Y. Sun, X. Chen, F.L. Heppner, T. Wyss-Coray, T.C. Sudhof, and M. Wernig. 2022. Treatment of a genetic brain disease by CNS-wide microglia replacement. Sci Transl Med 14:eabl9945.

47. Simard, A.R., D. Soulet, G. Gowing, J.P. Julien, and S. Rivest. 2006. Bone marrow-derived microglia play a critical role in restricting senile plaque formation in Alzheimer’s disease. Neuron 49:489–502.

48. Solomon, B., R. Koppel, D. Frankel, and E. Hanan-Aharon. 1997. Disaggregation of Alzheimer beta-amyloid by site-directed mAb. Proc Natl Acad Sci U S A 94:4109–4112.

49. Spangenberg, E., P.L. Severson, L.A. Hohsfield, J. Crapser, J. Zhang, E.A. Burton, Y. Zhang, W. Spevak, J. Lin, N.Y. Phan, G. Habets, A. Rymar, G. Tsang, J. Walters, M. Nespi, P. Singh, S. Broome, P. Ibrahim, C. Zhang, G. Bollag, B.L. West, and K.N. Green. 2019. Sustained microglial depletion with CSF1R inhibitor impairs parenchymal plaque development in an Alzheimer’s disease model. Nat Commun 10:3758.

50. Sperling, R., S. Salloway, D.J. Brooks, D. Tampieri, J. Barakos, N.C. Fox, M. Raskind, M. Sabbagh, L.S. Honig, A.P. Porsteinsson, I. Lieberburg, H.M. Arrighi, K.A. Morris, Y. Lu, E. Liu, K.M. Gregg, H.R. Brashear, G.G. Kinney, R. Black, and M. Grundman. 2012. Amyloid-related imaging abnormalities in patients with Alzheimer’s disease treated with bapineuzumab: a retrospective analysis. Lancet Neurol 11:241–249.

51. Spiteri, A.G., C.L. Wishart, R. Pamphlett, G. Locatelli, and N.J.C. King. 2022. Microglia and monocytes in inflammatory CNS disease: integrating phenotype and function. Acta Neuropathol 143:179–224.

52. Styren, S.D., R.L. Hamilton, G.C. Styren, and W.E. Klunk. 2000. X-34, a fluorescent derivative of Congo red: a novel histochemical stain for Alzheimer’s disease pathology. J Histochem Cytochem 48:1223–1232.

53. Tabata, Y., and Y. Ikada. 1988. Effect of the size and surface charge of polymer microspheres on their phagocytosis by macrophage. Biomaterials 9:356–362.

54. Tap, W.D., H. Gelderblom, E. Palmerini, J. Desai, S. Bauer, J.Y. Blay, T. Alcindor, K. Ganjoo, J. Martin-Broto, C.W. Ryan, D.M. Thomas, C. Peterfy, J.H. Healey, M. van de Sande, H.L. Gelhorn, D.E. Shuster, Q. Wang, A. Yver, H.H. Hsu, P.S. Lin, S. Tong-Starksen, S. Stacchiotti, A.J. Wagner, and E. investigators. 2019. Pexidartinib versus placebo for advanced tenosynovial giant cell tumour (ENLIVEN): a randomised phase 3 trial. Lancet 394:478–487.

55. van Dyck, C.H., C.J. Swanson, P. Aisen, R.J. Bateman, C. Chen, M. Gee, M. Kanekiyo, D. Li, L. Reyderman, S. Cohen, L. Froelich, S. Katayama, M. Sabbagh, B. Vellas, D. Watson, S. Dhadda, M. Irizarry, L.D. Kramer, and T. Iwatsubo. 2023. Lecanemab in Early Alzheimer’s Disease. N Engl J Med 388:9–21.

56. Wang, M., J. Munoz, A. Goy, F.L. Locke, C.A. Jacobson, B.T. Hill, J.M. Timmerman, H. Holmes, S. Jaglowski, I.W. Flinn, P.A. McSweeney, D.B. Miklos, J.M. Pagel, M.J. Kersten, N. Milpied, H. Fung, M.S. Topp, R. Houot, A. Beitinjaneh, W. Peng, L. Zheng, J.M. Rossi, R.K. Jain, A.V. Rao, and P.M. Reagan. 2020. KTE-X19 CAR T-Cell Therapy in Relapsed or Refractory Mantle-Cell Lymphoma. N Engl J Med 382:1331–1342.

57. Weller, R.O., D. Boche, and J.A. Nicoll. 2009. Microvasculature changes and cerebral amyloid angiopathy in Alzheimer’s disease and their potential impact on therapy. Acta Neuropathol 118:87–102.

58. Wilcock, D.M., A. Rojiani, A. Rosenthal, S. Subbarao, M.J. Freeman, M.N. Gordon, and D. Morgan. 2004. Passive immunotherapy against Abeta in aged APP-transgenic mice reverses cognitive deficits and depletes parenchymal amyloid deposits in spite of increased vascular amyloid and microhemorrhage. J Neuroinflammation 1:24.

59. Wilkinson, F.L., A. Sergijenko, K.J. Langford-Smith, M. Malinowska, R.F. Wynn, and B.W. Bigger. 2013. Busulfan conditioning enhances engraftment of hematopoietic donor-derived cells in the brain compared with irradiation. Mol Ther 21:868–876.

60. Xu, Z., Y. Rao, Y. Huang, T. Zhou, R. Feng, S. Xiong, T.F. Yuan, S. Qin, Y. Lu, X. Zhou, X. Li, B. Qin, Y. Mao, and B. Peng. 2020. Efficient Strategies for Microglia Replacement in the Central Nervous System. Cell Rep 32:108041.

61. Yan, P., K.W. Kim, Q. Xiao, X. Ma, L.R. Czerniewski, H. Liu, D.R. Rawnsley, Y. Yan, G.J. Randolph, S. Epelman, J.M. Lee, and A. Diwan. 2022. Peripheral monocyte-derived cells counter amyloid plaque pathogenesis in a mouse model of Alzheimer’s disease. J Clin Invest 132:

62. Youshani, A.S., S. Rowlston, C. O’Leary, G. Forte, H. Parker, A. Liao, B. Telfer, K. Williams, I.D. Kamaly-Asl, and B.W. Bigger. 2019. Non-myeloablative busulfan chimeric mouse models are less pro-inflammatory than head-shielded irradiation for studying immune cell interactions in brain tumours. J Neuroinflammation 16:25.

63. Yu, K., A.S. Youshani, F.L. Wilkinson, C. O’Leary, P. Cook, L. Laaniste, A. Liao, D. Mosses, C. Waugh, H. Shorrock, O. Pathmanaban, A. Macdonald, I. Kamaly-Asl, F. Roncaroli, and B.W. Bigger. 2019. A nonmyeloablative chimeric mouse model accurately defines microglia and macrophage contribution in glioma. Neuropathol Appl Neurobiol 45:119–140.

